# Characteristics of motor evoked potentials in patients with peripheral vascular disease

**DOI:** 10.1101/2023.08.10.552742

**Authors:** Pawandeep Sarai, Charlotte Luff, Cyrus Rohani-Shukla, Paul H Strutton

**Affiliations:** MSk Lab, Department of Surgery and Cancer, Division of Medicine, Imperial College London, London, United Kingdom

## Abstract

With an aging population, it is common to encounter people diagnosed with peripheral vascular disease (PVD). Some will undergo surgeries during which the spinal cord may be compromised and intraoperative neuromonitoring with motor evoked potentials (MEPs) is employed to help mitigate paralysis. No data exists on characteristics of MEPs in older, PVD patients, which would be valuable for patients undergoing spinal cord at-risk surgery or participating in neurophysiological research.

Transcranial magnetic stimulation, which can be delivered to the awake patient, was used to stimulate the motor cortex of 20 patients (mean (±SD) age 63.2yrs (±11.5) with confirmed PVD, every 10 minutes for one hour with MEPs recorded from selected upper and lower limb muscles. Data were compared to that from 20 healthy volunteers recruited for a protocol development study (28yrs (±7.6)). MEPs did not differ between patient’s symptomatic and non-symptomatic legs. MEP amplitudes were smaller in patients than in healthy participants in the upper limbs muscles, but not in lower limb muscles. Disease severity did not correlate with MEP amplitude. There were no differences over time in the coefficient of variation of MEP amplitude at each time point nor over the paradigm between groups. Latencies of MEPs were longer in patients for brachioradialis and vastus lateralis, but not in the other muscles studied.

The results obtained suggest PVD alone does not impact MEPs; there were no differences between more symptomatic and less symptomatic legs. Further, disease severity did not corelate with MEP characteristics. Differences observed in MEPs between patients and healthy participants are more likely a result of ageing.

With an aging population, more patients with PVD and cardiovascular risk factors will be participating in neurophysiological studies or undergoing surgery where spinal cord integrity is monitored. Our data show that MEPs from these patients can be easily evoked and interpreted.

## 1. Introduction

Motor evoked potentials (MEPs) are used extensively as a research tool to explore neuromuscular function in health and disease. [1–3] Changes in characteristics of MEPs have been observed in many neurological and neuromuscular pathologies, such as stroke and muscular dystrophies [3,4] and for intra-operative neuromonitoring (IONM) to diagnose and mitigate spinal cord injury during spinal surgery or thoraco-abdominal aortic aneurysm (TAAA) repair. [5,6]

Patients undergoing spinal surgery and TAAA repair are often older and frequently have cardiovascular risk factors such as high blood pressure, hypercholesterolaemia and smoke. [7,8] As such, they frequently have concomitant peripheral vascular disease (PVD) in addition to their presenting pathology, where the arteries of the lower limbs are atherosclerotic, leading to occlusion and subsequent chronic limb ischaemia (CLI). [9] This in turn leads to tissue loss and damage of the limb nerves and muscles.

MEP characteristics have been exhaustively studied in heathy volunteers, and in specific spinal or neurological disease. [10–13] The MEPs of PVD patients, however, have not previously been examined. Understanding changes that occur due to CLI is not only beneficial for IONM, but for any research or clinical situation where MEPs need to be characterised in patients with cardiovascular risk factors or recognised PVD. With an aging population and increasing rates of obesity and cardiovascular disease, [14] the co-incidence of PVD will inevitably increase. Current standards of IONM utilise MEPs generated by transcranial electrical stimulation (TES). [15] However, TES in the awake participant can be prohibitively painful. Transcranial magnetic stimulation (TMS) is an alternative technique for generating MEPs, with the benefit of being pain-free, [16] thus is frequently employed in neurophysiology research. [17] Further, MEP changes seen pre- and intra-operatively with TMS, correlate with TES-induced MEP changes during surgery and clinically after surgery. [17,18] TMS is therefore suitable to characterise MEPs in non-anaesthetised PVD patients in the research and surgical settings.

The aim of this study was to determine the characteristics of TMS-induced MEPs from limb muscles in PVD patients, and compare these to data from participants without cardiovascular risk, or PVD, which formed part of a protocol development study.

To our knowledge, this not been previously studied and would provide essential baseline data for future neurophysiological study of patients with cardiovascular disease and risk factors, with possible clinical applications, including intraoperative and post-operative monitoring

## 2. Material & Methods

### 2.1 Participants

20 patients with PVD (17 males), mean (±SD) age of 63.2 (±11.5) years took part in the study. All patients were symptomatic with radiologically confirmed PVD under the care of a vascular surgical team. Data were also collected prior to this study, from 20 participants (8 females; 28 (±7.6) years of age) with no co-morbidities or cardiovascular risk factors. Recruitment occurred between 15^th^ November 2016 and 20^th^ November 2018.

A TMS safety questionnaire was undertaken to ensure appropriateness for inclusion. [19] The inclusion and additional PVD-specific exclusion criteria for patients are listed in Table 1.

**Table 1:** Patient specific inclusion and exclusion criteria

| Inclusion Criteria | Exclusion Criteria |
| --- | --- |
| Experience symptomatic lower limb claudication pain (cramping in lower leg when walking), not currently suitable for surgery | Age < 18yrs |
| Under regular review by the Vascular Surgery Department at Imperial College NHS Trust for peripheral vascular disease (the build-up of fatty deposits in the arteries (blood vessels) that restrict blood supply in the lower legs) and not currently suitable for surgery. | Diabetes Mellitus (High blood sugar level) with no know microvascular complications e.g. peripheral neuropathy |
|  | Confirmed or possibility of pregnancy |
|  | Open wounds or skin lesions, including wet gangrene (dry gangrene permitted) |
|  | Inability to speak and understand English fluently |
|  | Patients with known thoracic or thoraco-abdominal aneurysm awaiting repair |

This study was approved by the Imperial College London research ethics committee (number: 16IC3553) and National Health Service health regulatory authority (number: 17/LO/1034) and performed with adherence to the Declaration of Helsinki guidance on conducting research in human participants. Written informed consent was obtained from all participants.

### 2.2. Disease Severity Questionnaire (VascuQol)

Patients completed the King’s College Hospital’s Vascular Quality of Life Questionnaire (VascuQol); this validated questionnaire determines disease severity with 25 questions exploring the effects of PVD on physical and emotional well-being of patients. [20] Each question is graded from 0 to 7, with 0 indicating PVD affects a particular aspect of their life all of the time (i.e. no ability to perform task). A mean score of 4.5 or less is associated with symptomatic intermittent claudication due to CLI. [21]

### 2.3. Electromyography

With participants supine, pairs of disposable Ag/AgCl electrodes (25mm diameter, 1041PTS, Henleys Medical Supplies Ltd., UK) were applied to the skin 20mm apart, overlying the muscles of interest, after skin preparation with alcohol. [22] To account for potential differences in disease severity between the left and right leg, patients had electrodes placed over left and right vastus lateralis (VL), tibialis anterior (TA) and abductor hallucis (AH), as well as left brachioradialis (BR) and left abductor pollicis brevis (APB)(Figure 1). A ground electrode was placed over the ulna olecranon process. These limb muscles were chosen as they are supplied by different major peripheral nerves.

**Figure 1:**
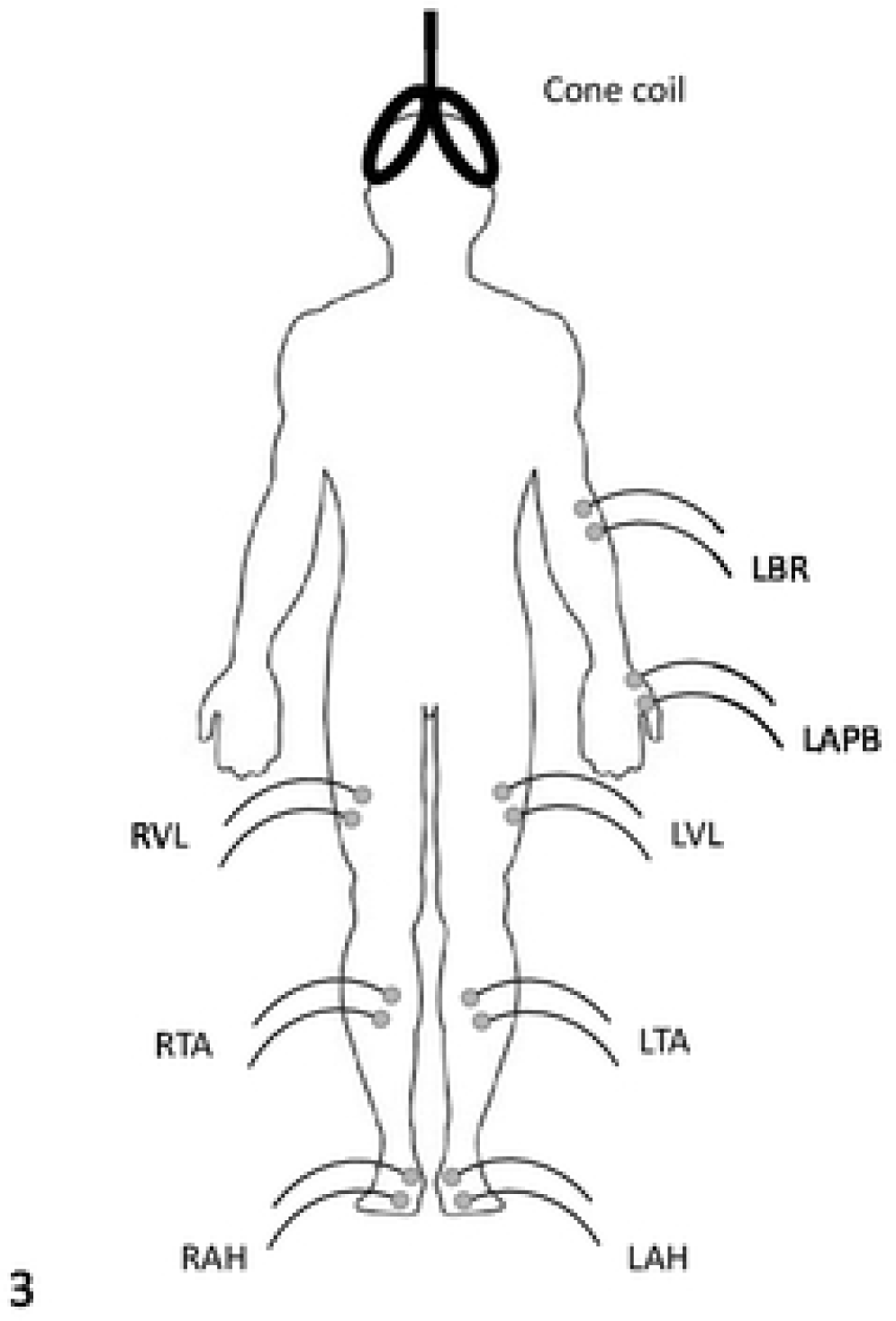
Electrode placement on PVD patients. Cone coil shown for illustration. Ground electrode over olecranon process omitted. L=Left, R=Right, BR = brachioradialis, APB = abductor pollicis brevis, VL = vastus lateralis, TA = tibialis anterior and AH = abductor hallucis.

Raw EMG signals were amplified (x1000), band-pass filtered (10Hz–1kHz) (ISO Dam, WPI, UK) and sampled (at 2kHz) using an analogue to digital converter (Power 1401, Cambridge Electronic Design [CED], UK). Signal software (CED, UK) was used to record to disk.

### 2.4. Transcranial Magnetic Stimulation

TMS was delivered using two Magstim 200² magnetic stimulators (Magstim, Dyfed, UK), each connected to a different coil, which was positioned over the vertex. A circular coil (90mm outer diameter) was used to stimulate upper limb muscles and was positioned B-side up to induce anticlockwise current in the brain, thus preferentially activating the right hemisphere. [23] An angled double-cone coil (wing outer diameter 120mm) was used to stimulate lower limb muscles and was positioned so that the induced current in the brain flowed in the posterior-to-anterior direction. A ‘combined’ resting motor threshold (cRMT) was found for the upper and lower limb muscles for each participant. This was the lowest stimulation intensity that produced MEPs ≥50µV in amplitude [24] in at least three out of six stimuli in either all upper or lower limb muscles, with the circular and double-cone coils, respectively. Intensities of 120% of the cRMT for the muscle with the highest threshold were used. [25]

Each block of stimulation consisted of six single stimuli from each coil, every 10 minutes over the course of 60 minutes (seven sets of twelve stimuli in total from 0 to 60 minutes). Participants were instructed to relax all muscles during stimulation to prevent facilitation. [26]

### 2.5. Data analysis

#### 2.5.1. Patient Disease Severity Questionnaire (VascuQol)

The scores from 25 questions were averaged to give a mean value from 0 to 7. Microsoft Excel 2016 (Microsoft, USA) was used to perform calculations.

#### 2.5.2. Motor Evoked Potentials

Signal Software was used for data analysis by visual positioning of cursors at the start and finish of each MEP. Peak-to-peak amplitude and latency (time from stimulation to first MEP deflection) were subsequently measured.

Given the known variability of TMS-induced MEPs in relaxed muscles [27] the variability both within a time point and across the whole experiment was calculated. The variability between measurements was determined by the coefficient of variation (CV); CV = standard deviation/mean.

##### 2.5.2.1 Variability within a time point

To determine the variability at a given time point, the amplitude of each MEP was measured, and the CV of the 6 MEPs calculated. The CV of the amplitude was calculated for each time point and for each muscle.

##### 2.5.2.2. Variability across the whole experiment

The averaged MEP (avMEP) is a digital averaging process, where the 6 MEPs at a given time point are amalgamated to produce a single MEP; from this the amplitude and latency are be measured.

To determine the variability of MEPs across the duration of the experiment, the amplitude of the avMEP at each of the 10-minute time points was measured and the CV calculated. This was performed for each muscle and each participant to calculate the mean CV of the avMEP amplitude; comparisons of the different upper and lower limb muscles were then performed.

### 2.6. Statistical analysis

Statistical analyses were carried out using Jamovi (The Jamovi project (2022), Jamovi (Version 2.3), Sydney, Australia). Data were tested for normality (Shapiro-Wilk) and log transformed where required. Parametric (t-tests and repeated measures ANOVA) or non-parametric (Mann-Whitney U-test, or Friedman test) were used as appropriate. Post-hoc multiple comparison procedures were performed to isolate the differences where a main effect was found.

AvMEP amplitudes, CVs of the avMEP amplitude, AvMEP latencies and CVs of the avMEP latency were collapsed over the hour and (where appropriate) compared between left and right sides in patients using paired t-tests.

As there were no differences between sides, data were averaged between sides and compared between patients and healthy participants using unpaired t-tests.

To examine changes in variability of MEP amplitudes within a time point over the hour protocol, CV of MEP amplitudes were compared over time and between left and right lower limb muscles using two-way repeated measures ANOVA (factors TIME and SIDE). Since no effect was seen for either time or side, data from left and right muscles were averaged. Two-way repeated measures ANOVA were used to assess for differences in CVs of MEP amplitudes over time and between patients and controls. Variability in amplitude and latency, CV of avMEP amplitude and CV of avMEP latency, respectively, across the 1-hour protocol for each muscle were compared using one-way repeated measures ANOVA. To determine the relationship between clinical score (VascuQol) and MEP characteristics, Spearman’s correlations were performed.

The level of statistical significance was set at P<0.05, or appropriate for multiple comparisons. Data are presented as mean ± SEM in the text and figures, unless stated otherwise.

## 3. Results

### 3.1 VascuQol Score in PVD Group

The VascuQol score was 4.3±1.2, representing symptomatic PVD not requiring surgical intervention (as per the main inclusion criteria). This score indicates patients experience cramp-like pain in the lower leg when walking due to muscle ischaemia.

### 3.2 Amplitude of avMEP

There were no differences in avMEP amplitudes between patient left and right legs (VL, left 0.34±0.10mV vs right 0.24±0.04mV, P=0.28; TA, left 0.63±0.14mV vs right 0.57±0.06mV, P=0.69; AH, left 0.80±0.13mV vs right 0.68±0.08mV, P=0.56). Data were therefore averaged between sides.

One-way RM ANOVA between averaged lower limb avMEP amplitudes (VL 0.29±0.06mV, TA 0.60±0.10mV and AH 0.74±0.10mV) and upper limb avMEP amplitudes (BR 0.22±0.05mV, APB 0.68±0.17mV), showed significant differences (F = 18.4, P<0.001). Post-hoc tests revealed AH was larger than VL (P<0.001) and BR (P<0.001); TA was larger than VL (P<0.001) and BR (P<0.001); and APB was larger than BR (P<0.001) and VL (P=0.048).

The avMEP amplitudes of BR and APB were smaller in patients (Mann-Whitney U-test) than in healthy participants (BR ([P] 0.22±0.05mV vs [H] 0.43±0.04mV, P<0.001; APB [P] 0.68±0.17mV vs [H] 1.87±0.46mV, P=0.013).

When collapsed across sides, there were no differences between the MEP amplitudes in the lower limb muscles of patients [P] and healthy [H] participants (VL [P] 0.29±0.06mV vs [H] 0.33±0.06mV, P=0.48; TA [P] 0.60±0.10mV vs [H] 0.67±0.08mV, P=0.43; AH [P] 0.74±0.10mV vs [H] 1.00±0.16mV, P=0.30).

### 3.3. CV of avMEP amplitude

There were no differences in CV of avMEP amplitudes between left and right legs in patients (VL, left 0.29±0.04 vs right 0.36±0.04, P=0.22; TA, left 0.30±0.04 vs right 0.30±0.04, P=0.69; AH, left 0.23±0.02 vs right 0.20±0.02, P=0.25). Data were therefore averaged between sides.

One-way RM ANOVA (Friedman) between averaged lower limb CV of avMEP amplitudes (VL 0.33±0.03, TA 0.30±0.03 and AH 0.22±0.02) and upper limb CV of avMEP amplitudes (BR 0.28±0.02, APB 0.50±0.04), showed a significant difference (F = 12.8 P<0.001, see Figure 2). CV of avMEP for APB was larger than BR (P=0.001), AH (P<0.001), TA (P=0.013) and VL (P=0.017); and CV of avMEP for AH was smaller than VL (P=0.019) and TA (P=0.017).

**Figure 2:**
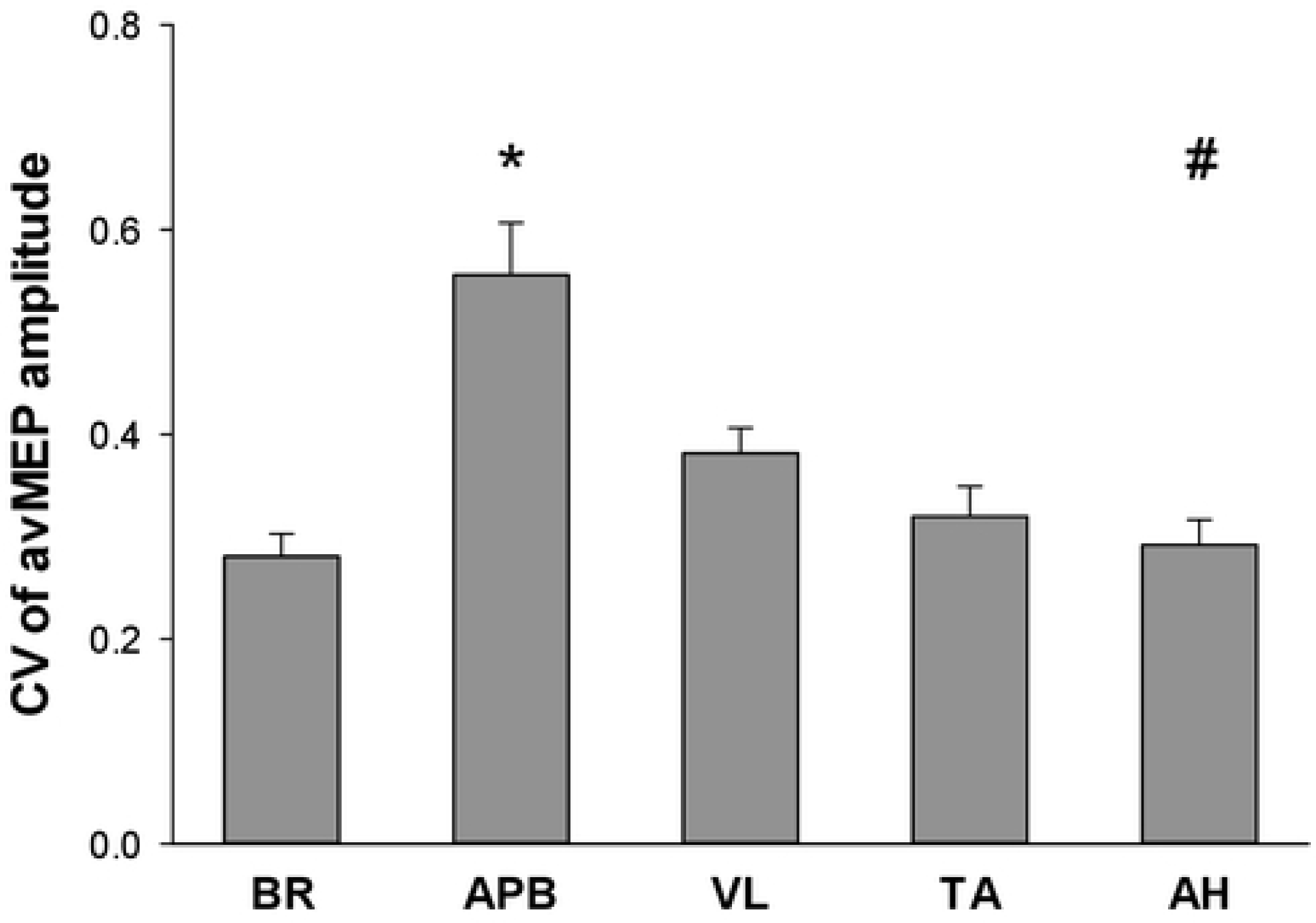
CV of avMEP amplitude in PVD patients (mean+SEM). BR – brachioradialis, APB – abductor pollicis brevis, VL – vastus lateralis, TA – tibialis anterior and AH – abductor hallucis. * denotes a significant difference between APB and all other muscles (P<0.05). # denotes a significant difference between AH and VL and between AH and TA (P<0.05).

When compared to health participants, RM ANOVA revealed an effect of muscle (F= 9.44, P<0.001), but no effect of group (F=23.8, P=0.13) and no interaction (F=2.06, P=0.09) Post-hoc analysis showed ABP was greater than BR (P=0.04), AH (P<0.001) and TA (P=0.02), BR was greater than AH (P=0.007); and VL was greater than AH (P<0.001).

### 3.4 Latency of avMEP

There were no differences in avMEP latencies between left and right legs in patients (VL, left 25.46±0.71ms vs right 25.58±0.68ms, P=0.32; TA, left 30.96±0.96ms vs right 31.71±0.82ms, P=0.17; AH, left 44.39±1.31ms vs right 43.76±1.45ms, P=0.22). Data were therefore averaged between sides.

Log transformation did not normally distribute the data. One-way RM ANOVA (Friedman) between averaged lower limb avMEP latency (VL 25.61±0.65ms, TA 31.33±0.85ms and AH 44.08±1.36ms) and upper limb avMEP latency (BR 17.21±0.31ms and APB 23.79±0.52ms), showed significant differences (χ^2^=72.5, P<0.001). All avMEP latencies were different between patient muscles (P <0.001).

MEP latencies were longer in patients compared to healthy participants (Mann-Whitney U-test) in the upper limb muscles, (BR [P]: 17.21±0.31ms vs [H] 14.84±0.46ms, P<0.001; APB [P]: 23.79±0.52ms vs [H] 21.27±0.55ms, P<0.001).

When collapsed across sides, the avMEP latencies in the lower limb muscles of patients were longer than healthy participants for VL ([P]: 25.61±0.65ms vs [H] 24.18±0.88ms, P=0.049), TA ([P] 31.33±0.85ms vs [H] 29.37±0.57ms, P=0.007), and AH ([P] 44.08±1.36ms vs [H] 39.62±0.65, P<0.001).

### 3.5. CV of avMEP latency

The CV avMEP latency between patient’s left and right legs demonstrated no differences (VL, left 0.04±0.01 vs right 0.03±0.01, P=0.21; TA, left 0.04±0.01 vs right 0.03±0.004, P=0.95; AH, left 0.03±0.01 vs right 0.02±0.004, P=0.04). Data were therefore averaged between sides.

One-way RM ANOVA (Friedman) between averaged lower limb CV avMEP latency (VL 0.04±0.01, TA 0.03±0.003 and AH 0.02±0.002) and upper limb CV avMEP latency (BR 0.06±0.006, APB 0.04±0.06), showed significant differences (χ^2^=21.5, P<0.001). AH was smaller than VL (P=0.044), TA (P=0.034), BR (P<0.001) and APB (P=0.002); and BR was larger than TA (P=0.004) and VL (P=0.003; see Figure 3).

**Figure 3:**
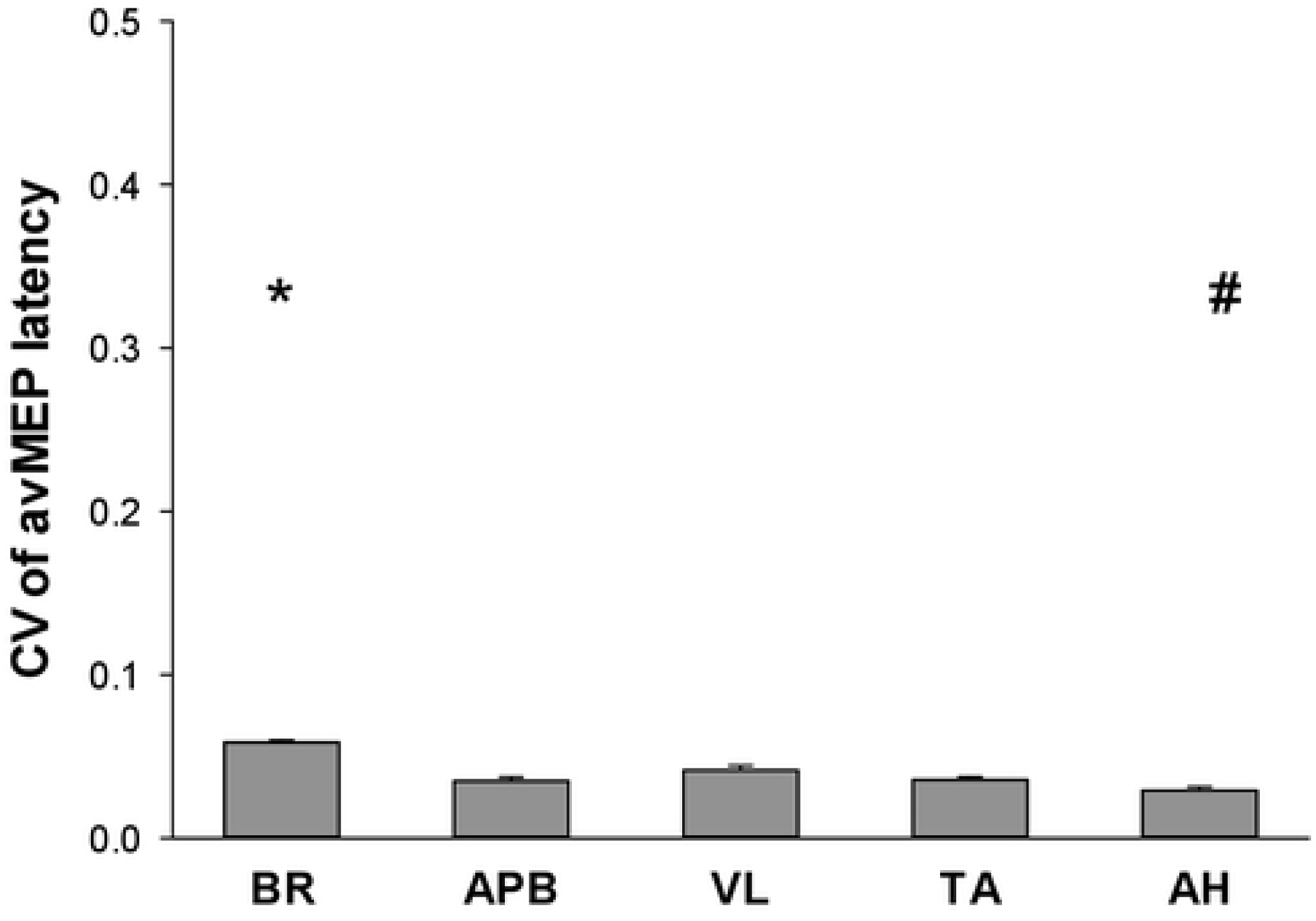
CV of avMEP latency for each patient muscle (mean+SEM). BR – brachioradialis, APB – abductor pollicis brevis, VL – vastus lateralis, TA – tibialis anterior and AH – abductor hallucis. * denotes a significant difference between BR and TA and between BR and VL (P<0.01); and # denotes a significant difference between AH and all other muscles (P<0.05).

There were no differences in CV of avMEP latency between patients and healthy participants (BR ([P]: 0.06±0.006 vs [H] 0.06±0.001, P=0.87); APB ([P]: 0.04±0.06 vs [H] 0.05±0.003, P=0.93).

Similarly, there were no differences between the CV of avMEP latency in the lower limb muscles of patients and healthy participants (VL [P]: 0.04±0.01 vs [H] 0.06±0.005, P=0.32; TA [P] 0.03±0.003 vs [H] 0.02±0.001, P=0.237; AH [P] 0.02±0.002 vs [H] 0.03±0.001, P=0.88).

### 3.6 Mean CV of MEP amplitude at each time point

Data were collapsed across sides as avMEP amplitude and CV of avMEP amplitude were not different between sides. ANOVA comparing the CV of MEP amplitude at each 10-minute time point revealed no effect of time for any muscles (F=0.0908, P=0.49) ANOVA revealed no effect of group on the CV of MEP amplitude between the patients and healthy participants (F=0.503, P =0.483) at each 10-minute time point (see Figure 4).

**Figure 4:**
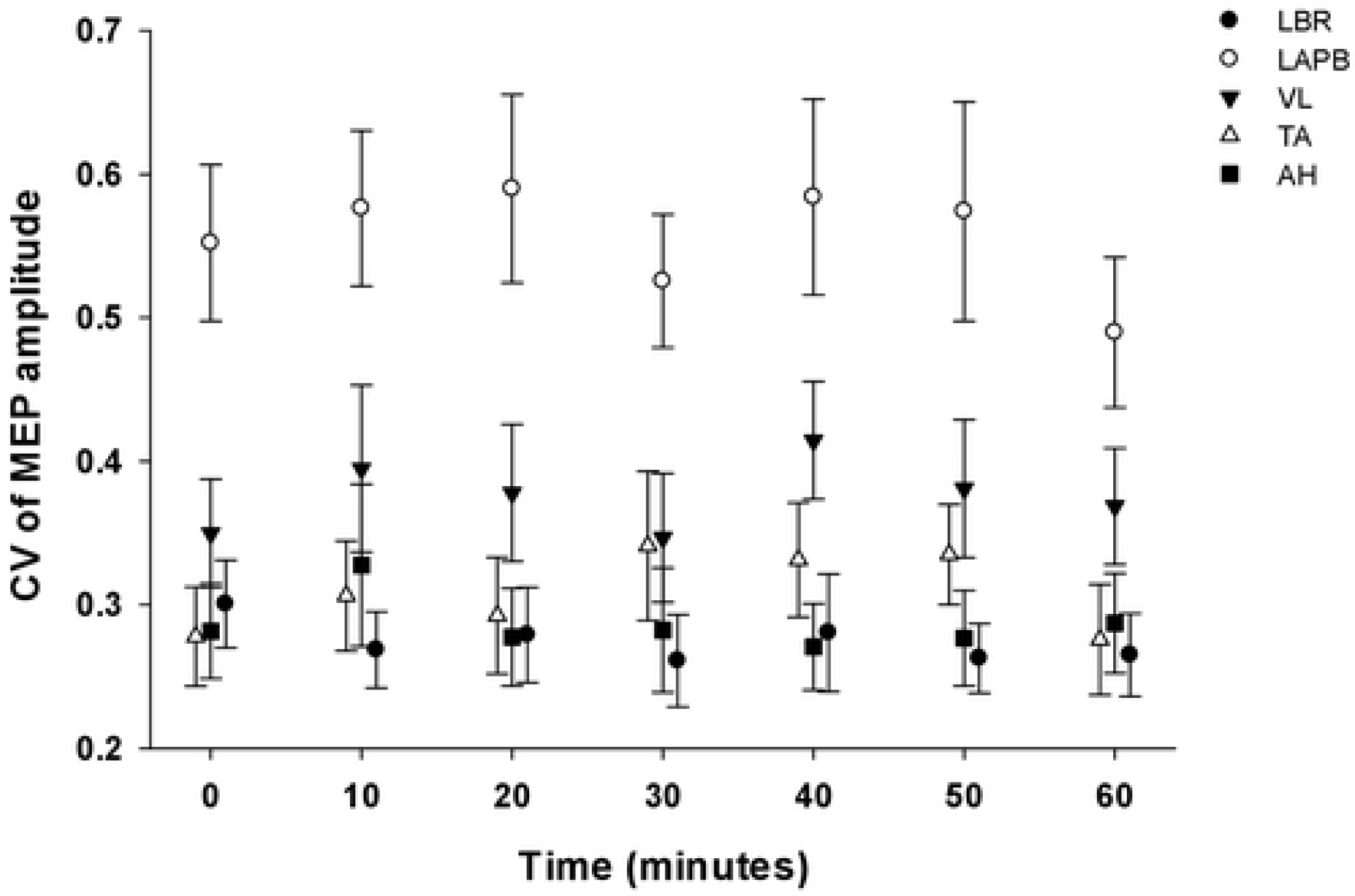
Coefficient of variation of avMEP amplitude (±SEM) of MEP amplitude for each muscle at each time point in PVD patients. Left brachioradialis (LBR), left abductor pollicis brevis (LAPB), vastus lateralis (VL), tibialis anterior (TA) and abductor hallucis (AH). Values for TA and LBR offset by -1 minute and +1 minute, respectively, for graphical clarity.

### 3.7 Correlations

#### 3.7.1 VascuQol score and MEP amplitudes

There was no correlation between VascuQol score and MEP amplitudes (ρ range -0.373-0.066, P>0.05 for all comparisons), for any muscle, except VL (for VL (ρ -0.781, P<0.01), For VL, disease severity score was inversely proportional to MEP amplitude, i.e. worse PVD was associated with larger MEP amplitudes.

#### 3.7.2 Mean CV of avMEP and mean avMEP amplitude

There was no correlation between the CV of the avMEP and the mean avMEP amplitude of any muscle (ρ range -0.440 to - 0.065, P>0.05 in all cases).

## 4 Discussion

The results of this study have demonstrated MEP characteristics of patients with PVD. When compared to our previous data (unpublished), they appear to be similar to healthy, younger participants, with differences likely attributable to age rather than disease. In the following, these findings will be discussed further, along with their potential impact on future neurophysiological research and clinical application.

### 4.1 MEP Amplitude and Latency

PVD patients have MEPs with characteristics consistent with those in the literature attributable to the effects of aging. [28,29] These studies commonly compare healthy younger participants with matched healthy older participants, and have found both at rest and during contraction, older participants have smaller MEP amplitudes with longer latencies.

The data from this investigation found upper limb muscles MEPs with significantly smaller amplitudes and longer latencies, when compared to healthy participant data from a previous methodological study undertaken by the research group. In the lower limb muscles, there were no differences in MEP amplitudes or latencies between the symptomatic (presumably more diseased) and asymptomatic legs, allowing the data for the left and right leg to be averaged. This further supports the negligible impact of the PVD on MEPs. Patient MEP amplitudes in VL, TA and AH showed a trend towards being smaller when compared to healthy data, although this was not significant. As with the upper limbs, lower limb MEP latencies were longer in patients than in healthy participants, in keeping with the age-related changes reported in the literature.

A single significant correlation was found between the disease severity score and avMEP amplitude; higher VascuQol scores were associated with smaller VL amplitude. The VascuQol is a functional scoring system, assessing the patient’s ability to perform common daily activities, such as bathing; a maximum score of 7 would indicate ‘maximum’ function with no impact on function due to PVD. Therefore, with more severe disease (i.e. a lower VascuQol score), one would predict smaller MEPs secondary to potential tissue damage. No such correlations were found, and the single negative correlation is difficult to explain. It is therefore proposed symptomatic PVD, which impairs activities of daily living, appears to have little impact on limb muscle MEP characteristics.

The protocol was designed to include patients who had stable disease with symptomatic leg pain on exercise, not severe enough to require surgery; the mean VascuQol score reflects this. These patients have an arterial blood supply to their lower limbs which is adequate provided no exertional stress is placed upon them; there is a state of intermittent ischaemia. Based on our results, this degree of ischaemia does not seem to result in any deterioration of nerve function. To the best of our knowledge, this has not been investigated before and provides valuable baseline data for older patients with known cardiovascular morbidity, with or without diagnosed PVD (given identical risk factors) undergoing neurophysiological investigation or surgery where IONM is employed.

We were aware of the potential effect of MEP amplitude on the variability of MEPs within a muscle. The design of our protocol meant there was a disparity between the relative stimulation intensity for each muscle given a single ‘combined’ test intensity per coil was used. A stimulus intensity of 120% of motor threshold of the muscle with the highest threshold was used; therefore, the intensity would be greater for the remaining muscles with lower motor thresholds. Previous work has shown greater stimulation intensities result in MEPs with not only larger amplitudes but also less variability, likely through consistent activation of most of the available motor units. [25,30,31] However, our results showed no correlation between the mean CV of avMEP and the mean amplitude of avMEP for any muscle, indicating negligible impact from using a single stimulation intensity at 120% of cRMT.

Amplitude is a key factor to take into consideration when considering the use of MEPs for clinical purposes. A sudden reduction in amplitude or loss of MEPs is highly suggestive of injury during spinal surgery. [32] Muscles with small MEP amplitudes can make it difficult to detect injury-induced decreases. Furthermore, the depressive effects anaesthetic agents significantly reduce MEP amplitudes and prolong latencies. [18,33] In many IONM guidelines, usually a hand and foot muscle are used as control and monitoring muscle, respectively. [34] The results of this study show that APB (hand) and AH (foot) in PVD patients produced the largest mean amplitudes of the upper and lower limb muscles, respectively, similarly observed in healthy participants; thus they would also be suitable for monitoring purposes in the PVD population as per standard IOMN guidelines. [34]

### 4.2 Reliability

Many neurophysiological studies have investigated the variability of TMS-induced MEPs. [10,27,30,35–37] No previous study to our knowledge has compared the variability of different muscles over a prolonged time period. As can be seen from the current data, it is evident that there is variability in TMS-induced MEP characteristics between stimuli at each epoch and over time. Our interest lay in the extent of this variation between muscles and compared to healthy participants.

During surgeries where the spinal cord is vulnerable, there will be certain surgical interventions or time points where there is a higher SCI risk. At these crucial timepoints, there will be a need for repeated trains of stimuli to continually assess spinal cord integrity, hence the need to assess variability within trains at a given time point (CV of mean MEP). Determining the variability of TMS-induced MEPs over time is also necessary particularly where frequent but intermittent monitoring is required for a longer duration of time, hence the need to assess variability over an extended period (CV of avMEP). This was achieved with a protocol recording trains of six MEPs at regular intervals over a 1-hour period.

TMS-induced MEPs have been shown to vary in amplitude; [31] this variability is greater with TMS than with TES, since motor cortical activation occurs largely trans-synaptically. [27,38] TES induces MEPs through direct activation of corticospinal neurons and generates more consistent results. [39] It is for this reason that surgical neuromonitoring guidelines usually recommend TES. [15] However, there are practical disadvantages to its use, such as pain, which limits its use in awake participant.

Our results have shown that the CVs of the mean avMEP amplitude are comparable in patients and healthy participants, as no effect of group was found. The mean CV of the avMEP amplitude gives an indication of the variation of all MEPs generated in each muscle over the course of the 1-hour protocol. This is especially true for BR and AH, which had the lowest CVs of the avMEP amplitude for upper and lower limb muscles in both groups of participants. These results are in keeping with previous studies, showing TMS-related measurements have good intra- and inter-investigator [35] and test-retest [37] reliabilities in healthy young and old participants. [10,31,40] Given there were no significant differences between healthy participants and PVD patient CV values in this study, the variability of normative MEP data in the literature where predominantly young, healthy participants are included, should be applicable to patients with cardiovascular disease or risk factors participating in research or neurophysiological testing.

Some of the variability in MEPs could be attributed to methodology, [41–43] such as stimulus intensity or number of stimuli delivered, [10] level of muscle contraction at time of stimulation, [44,45] data analysis procedures including removal of initial MEPs when averaging [30] and technical factors relating to coil position. [46,47]. To limit this, we delivered trains of 6 stimuli for our averaging calculations, maintained strict coil orientation and position, performed stimulation at rest (as confirmed by pre-stimulus EMG levels) and removed only data where there was muscle contraction. As a result, MEP measures in this investigation, including variability, are comparable to previously published work. [27,31,48]

### 4.3 Limitations

The authors acknowledge the comparison with data from young healthy volunteers, rather than healthy age-matched participants, may be viewed as a limitation. The healthy volunteer data were generated from a protocol development study, undertaken before the current study. The comparison presented here is to demonstrate differences between diseased and disease-free participants. PVD is a disease of aging and associated with lifestyle factors, with the atherosclerotic process increasing with age; age is an independent risk factor, irrespective of health status. [49] Thus, it would be difficult to find an older person who would be truly disease-free and “healthy” with respect to their arterial wall integrity. Confirmation would require advanced functional assessments and blood flow imaging studies, [50] which would have been practically prohibitive for this study. Data exist for healthy older participants and our data are compared to the literature in the discussion.

The use of a cRMT results in a stimulation intensity of 120% for the muscle with the highest motor threshold, but greater for the other muscles. It is convention to use a specific coil and test intensity to target each muscle but would have been impractical in the confines of a time critical protocol. However, previous work has shown only small differences in motor threshold between limb muscles, [3,51] thus the use of a single test intensity is unlikely to have a major impact on the data generated.

In many neurophysiological studies, MEP amplitudes are often normalised to the maximum M-waves (MMax), the maximal EMG response to direct motor nerve stimulation; [52] this allows comparisons to be made across muscles and between participants with different MEP amplitudes. However, since a key focus of the current investigation was variability of MEPs over time (as assessed by CV), normalisation to MMax was not demmed necessary and due to the time critical nature of this protocol, it was not feasible to include measurement of MMax for all muscles studies.

### 4.4 Conclusions & Recommendations

MEPs were recordable from upper and lower limb muscles in PVD patients and were found to have similar characteristics to data from our healthy participants. Our results suggest there are no additional effects of pathological limb ischaemia on MEP characteristics beyond those expected by aging. With an increasingly aging population, there will be greater numbers of patients with PVD and cardiovascular risk factors participating in neurophysiological studies or undergoing surgery where spinal cord integrity is monitored [53] and MEPs from these patients can be easily evoked and interpreted.

